# Cas12a trans-cleavage can be modulated in vitro and is active on ssDNA, dsDNA, and RNA

**DOI:** 10.1101/600890

**Authors:** Ryan T. Fuchs, Jennifer Curcuru, Megumu Mabuchi, Paul Yourik, G. Brett Robb

## Abstract

CRISPR-Cas12a (Cpf1) are RNA-guided nuclease effectors of acquired immune response that act in their native organisms by cleaving targeted DNA sequences. Like CRISPR-Cas9 RNA-guided DNA targeting enzymes, Cas12a orthologs have been repurposed for genome editing in non-native organisms and for DNA manipulation *in vitro*. Recent studies have shown that activation of Cas12a via guide RNA-target DNA pairing causes multiple turnover, non-specific ssDNA degradation in *trans*, after single turnover on-target cleavage in *cis*. We find that the non-specific *trans* nuclease activity affects RNA and dsDNA in addition to ssDNA, an activity made more evident by adjustment of reaction buffer composition. The magnitude of the *trans* nuclease activity varies depending on features of the guide RNA being used, specifically target sequence composition and length. We also find that the magnitude of *trans* nuclease activity varies between the three most well-studied Cas12a orthologs and that the Cas12a from *Lachnospiraceae* bacterium ND2006 appears to be the most active.

## INTRODUCTION

Clustered Regularly Interspaced Short Palindromic Repeats (CRISPR) are DNA sequences associated with acquired immune response systems in bacteria and archaea (1–3). There is extensive diversity amongst the known CRISPR systems, but a common characteristic is that they use a targeted effector protein or protein complex to cleave foreign nucleic acid using either an RNA or DNA guide (4,5). The effector proteins Cas9 and Cas12a (previously known as Cpf1) from Class 2 Type II and Type V-A CRISPR systems, respectively, use RNA guides to cleave dsDNA in a targeted manner and have been repurposed for genome editing applications in eukaryotic and prokaryotic organisms (6–15).

While both Cas9 and Cas12a have been used for similar applications, they are evolutionarily distinct proteins and achieve specific DNA targeting by different mechanisms. Key properties of Cas9 include that it utilizes dual RNA molecules for targeting – known as crRNA and tracrRNA – which are processed by other proteins, requires a protospacer adjacent motif (PAM) in the target DNA proximal to the cleavage site, uses a RuvC domain to cleave the non-target DNA strand and an HNH-domain to cleave the target DNA strand, and typically leaves blunt ends on the cleavage products which it remains bound to post cleavage (16–19). Additionally for Cas9, it has been shown that crRNA and tracrRNA can be fused into a ~100 nucleotide (nt) single-guide (sgRNA) and that the 3’ end of the non-target strand can be partially degraded after the blunt cut is made to leave a 5’ overhang on the PAM-distal side of the cleavage site (16,20). In contrast, Cas12a processes its own ~40 nt guide RNAs (crRNA) from primary transcripts, requires a T-rich PAM that is distal to the cleavage site, uses a single RuvC domain to sequentially cleave each strand of the DNA target, typically leaves 5’ overhangs on the cleavage products, and remains bound to the PAM-proximal cleaved DNA while the PAM-distal piece of DNA dissociates (11,21–26). Recent work has identified an additional activity attributed to Cas12a, specifically that Cas12a ribonucleoprotein (RNP) remains in an activated state by remaining bound to the crRNA and PAM proximal DNA after release of the cleaved, PAM distal DNA. This activated state leads to non-specific degradation of ssDNA in *trans* that is attributed to the RuvC domain (27–29).

In this work we investigate the *trans* nuclease activity of Cas12a and find that while ssDNA is the most susceptible substrate, the *trans* activity also results in degradation of RNA and dsDNA as there has been some evidence of in other studies (27,28). The degradation of RNA and dsDNA can either be nearly imperceptible over typical reaction timescales or can be extensive depending on reaction buffer composition. In addition, we find that *trans* nuclease activity does not appear to be influenced by non-target DNA sequence, but guide RNA parameters can alter the relative amount of the activity without significantly affecting the yield of on-target *cis* cleavage over the course of typical reaction lengths. We show that the three most well-studied orthologs of Cas12a all have *trans* nuclease activity, but the magnitude in identical reaction conditions varies significantly among orthologs and the Cas12a from *Lachnospiraceae* bacterium ND2006 (LbaCas12a) repeatedly showing the most predominant *trans* activity. Future work will be needed to fully understand the ramifications of *trans* activity for both *in vitro* and *in vivo* applications of Cas12a.

## MATERIALS AND METHODS

### Oligonucleotides

All DNA and RNA oligonucleotides were synthesized by Integrated DNA Technologies (IDT, Coralville IA) or MilliporeSigma (St. Louis, MO) and the sequences are available in Supplementary Table S1. Mature crRNA sequences for Lba, Asp, and FnoCas12a were obtained from a previous publication and were appended with either a 20 nt or 24 nt targeting sequence (11).

### Cas12a purification and guide RNA design

EnGen^®^ LbaCas12a from New England Biolabs (NEB #M0653, Ipswich, MA) was used for all experiments involving LbaCas12a. AspCas12a was obtained from Integrated DNA Technologies (IDT; Alt-R^®^ A.s. Cas12a (Cpf1) V3). The Cas12a from *Francisella tularensis* subsp. *novicida* strain U112 (FnoCas12a) was purified via the following scheme. Recombinant protein was expressed in modified *E. coli* NiCo21 (DE3) cells (NEB #C2925) harboring the Cas12a ortholog expression plasmid by growing in LB at 37°C followed by induction of expression at 23°C for 16 hr in presence of 0.4 mM IPTG. Cells were disrupted by sonication prior to chromatographic purification. Recombinant protein was purified using HiTrap DEAE^FF^ (GE Healthcare, Chicago IL), HisTrap^HP^ (Ni-NTA) (GE Healthcare) and HiTrapSP ^HP^ (GE Healthcare) columns and were dialyzed and concentrated into 20 mM Tris-HCl (pH 7.4), 500 mM NaCl, 1 mM DTT, 0.1 mM EDTA and 50% glycerol.

### Cas12a *in vitro* digests

Cas12a guide RNAs were heated to 65°C, slow cooled to room temperature and quantified by Nanodrop 2000 (Thermo-Fisher, Waltham MA) analysis prior to being used as a stock for *in vitro* digest experiments. All Cas12a RNP’s were formed by mixing crRNA and protein in a 1.1X buffer at 25°C for 10 minutes. DNA addition reduced the buffer concentration to 1X and reactions were incubated at 37°C unless otherwise indicated for a variable amount of time depending on the experiment. Reactions were typically carried out in 1X NEBuffer 2.1 (50 mM NaCl, 10 mM Tris-HCl, 10 mM MgCl_2_, 100 ug/ml BSA, pH 7.9) unless otherwise indicated, with the exception of AspCas12a where an unique buffer was used (50 mM NaCl, 10 mM Tris-HCl, 10 mM MgCl_2_, 1 mM DTT, pH 6.5). Reactions were quenched by addition of 8-fold molar excess of EDTA relative to the Mg^2+^ present. Immediately following EDTA addition, DNA was purified by using a Monarch^®^ PCR&DNA cleanup kit (NEB #T1030) before analyzing the DNA by capillary electrophoresis (30), PAGE, or next generation sequencing. DNA substrates used for digestion reactions were either generated by PCR using Q5^®^ High-Fidelity 2X Hot Start Master Mix (NEB #M0494), oligonucleotides produced by IDT, or commercially available products such as M13mp18 single stranded DNA (NEB #N4040).

### Guide RNA Binding Assay

Synthesized guide RNAs were 5’-labeled with ^32^P using T4 PNK (NEB #M0201) and [γ-^32^P]ATP (Perkin Elmer #BLU002A250UC, Waltham MA) per manufacturer protocol and reaction products were purified with the Zymo Clean & Concentrator - 5 kit (Zymo Research #R1016, Irvine CA). RNA concentration post labeling was measured with the Qubit RNA HS kit (ThermoFisher #Q32852). EnGen^®^ LbaCas12a was titrated in the presence of 50 pM ^32^P-gRNA and reactions were incubated for 40 min at room temperature (25°C). 30 µL of each reaction was combined with 2 µL of loading dye (50% glycerol, 0.02% xylene cyanol) and resolved by immediately loading on a 6% Tris-Borate-EDTA acrylamide gel in the cold room. Gels were scanned with an Amersham™ Typhoon™ Biomolecular Imager (GE Healthcare) and imaged products were quantified with ImageQuant™ software. The fraction of shifted gRNA (gRNA-Cas12a ribonucleoprotein complex) was plotted versus the concentration of Cas12a protein in the reaction and fit with a standard hyperbolic binding isotherm using the KaleidaGraph software (31). All experiments were carried out at least 3 times.

### Next generation sequencing

Libraries for next generation sequencing were prepared by using the NEBNext^®^ Ultra™ II DNA Library Prep Kit for Illumina^®^ (NEB #E7645) following the protocol supplied with the kit and individual samples were indexed during the PCR step with NEBNext^®^ Multiplex Oligos for Illumina^®^. The resulting libraries were analyzed using an Agilent 2100 Bioanalyzer (Agilent Technologies, Santa Clara CA), pooled and submitted for sequencing on either an Illumina^®^ Miseq or Nextseq instrument. Sequencing data was analyzed and mapped to known reference sequences using standard tools in a local instance of Galaxy. For the data in Figure 2B, sequence coverage at each position was normalized by dividing by the value for the position with the highest coverage. For Figure 2C, coverage at positions 1-19 in the randomized region of DNA was normalized for each sample by dividing by the coverage at position 20. Sequence logos were made by loading a Position Weight Matrix into the enoLOGOS web browser tool (32). Position weight for each base at each position was calculated by the following equation:

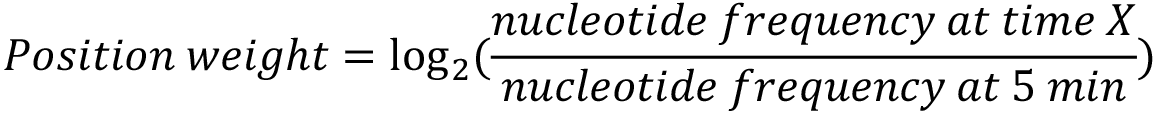

### Fluorescent reporter assay for *trans* nuclease activity

All Cas12a RNPs were formed as described above (Cas12a *in vitro* digests). Enzyme and crRNA were incubated at a 1:1 molar ratio and a final concentration of 250 nM. A 103 bp DNA containing the target sequence (activator) was added to the assembled RNPs at a final concentration of 25 nM, and incubated at 37°C for 30 minutes to allow cleavage of the target DNA. For assessment of *trans* activity of the activated RNPs, 50 pmol of ssDNA reporter oligo (5’-FAM-NNNNN-Q-3’) or ssRNA reporter (5’-FAM-rNrNrNrNrN-Q-3’) mixed with the RNPs in a black 96-well plate (Costar #3615). Recording of fluorescence intensity was started immediately via a SpectraMax plate reader. Data was gathered in 15 second intervals for either 1 hour (ssDNA) or 3 hours (ssRNA), with the exception of the data collected for the ortholog *trans* activity. In this case, data was collected for either 2 hours (ssDNA) or 16 hours (ssRNA). The data were fit with a single exponential equation in Kaleidograph software as follows:

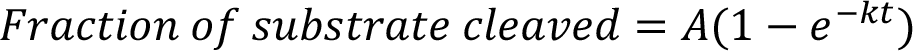

A is amplitude, k is the observed rate constant (k_obs_), and t is time.

## RESULTS

### LbaCas12a has *trans* nuclease activity that acts on ssDNA and dsDNA

We first observed Cas12a *trans* nuclease activity in *in vitro* cleavage assays using LbaCas12a and 5’-FAM-labeled DNA substrates where reactions products were analyzed by capillary electrophoresis. When incubated for longer than 5 min, reaction products routinely ran either as a smear or the FAM label migrated in a position consistent with its removal from the DNA (data not shown). While in the process of characterizing this activity, the first reports describing the *trans* activity of Cas12a were published (27,28). We investigated the activity in additional detail. We first tested the ability of LbaCas12a coupled to a guide RNA complementary to a small, linear dsDNA target to degrade a non-target, circular ssDNA in *trans*. As shown in Figure 1A, the 7.2 kilobase (kb) circular ssDNA was degraded rapidly by LbaCas12a such that no intact substrate was detected after 10 seconds and degraded products were no longer visible after 5 minutes. Next, we wondered whether other DNA species such as linear dsDNA or circular supercoiled dsDNA (pUC19) could be affected by *trans* activity. Figure 1B shows that over time pUC19 was converted into a mixture of nicked and linearized species when LbaCas12a was activated by a guide RNA and target DNA, but in the absence of target DNA the pUC19 DNA remained supercoiled. These results demonstrate that the *trans* nuclease activity acts as an endonuclease. A similar experiment was performed on a ladder of linear dsDNA products (Figure 1C). The dsDNA products were converted into a smear of degraded products when guide RNA and complementary target DNA were included, and the degradation increased over time. It is noteworthy that consistent with previous results (27,28) the most susceptible substrate for *trans* activity appears to be ssDNA as complete degradation occurs on the seconds to minutes timescale versus minutes to hours for dsDNA. To gain insight into the nature of the dsDNA fragments contained in the smear, we performed an *in vitro* cleavage experiment with a 226 bp dsDNA and LbaCas12a loaded with a guide that targeted the DNA. Reaction products were sampled over time and subjected to Illumina sequencing (Figure 2A). We observed that the sequence coverage is reduced at the DNA ends (Figure 2B). Three of the four ends resulting from the two pieces of cleaved DNA were affected. The end that contained the target sequence was unaffected. This observation is consistent with it remaining bound to the activated LbaCas12a RNP (24–26) and so it is not surprising that it was protected from degradation. The DNA library preparation method used has end repair and ligation steps, so this approach does not detect DNA nicks or overhangs thus providing an informative but incomplete picture of the degradation products. The fact that pUC19 covalently closed circular DNA is nicked by *trans* activity (Figure 1B) suggests that the linear dsDNA is being internally nicked and it is likely that accumulation of nicks at the end of the molecule causes dsDNA ends to fall apart and not be captured by the library construction method. Taken together, these observations show that both ssDNA and dsDNA are subject to degradation in *trans* by activated Cas12a RNPs.

**Figure 1.**
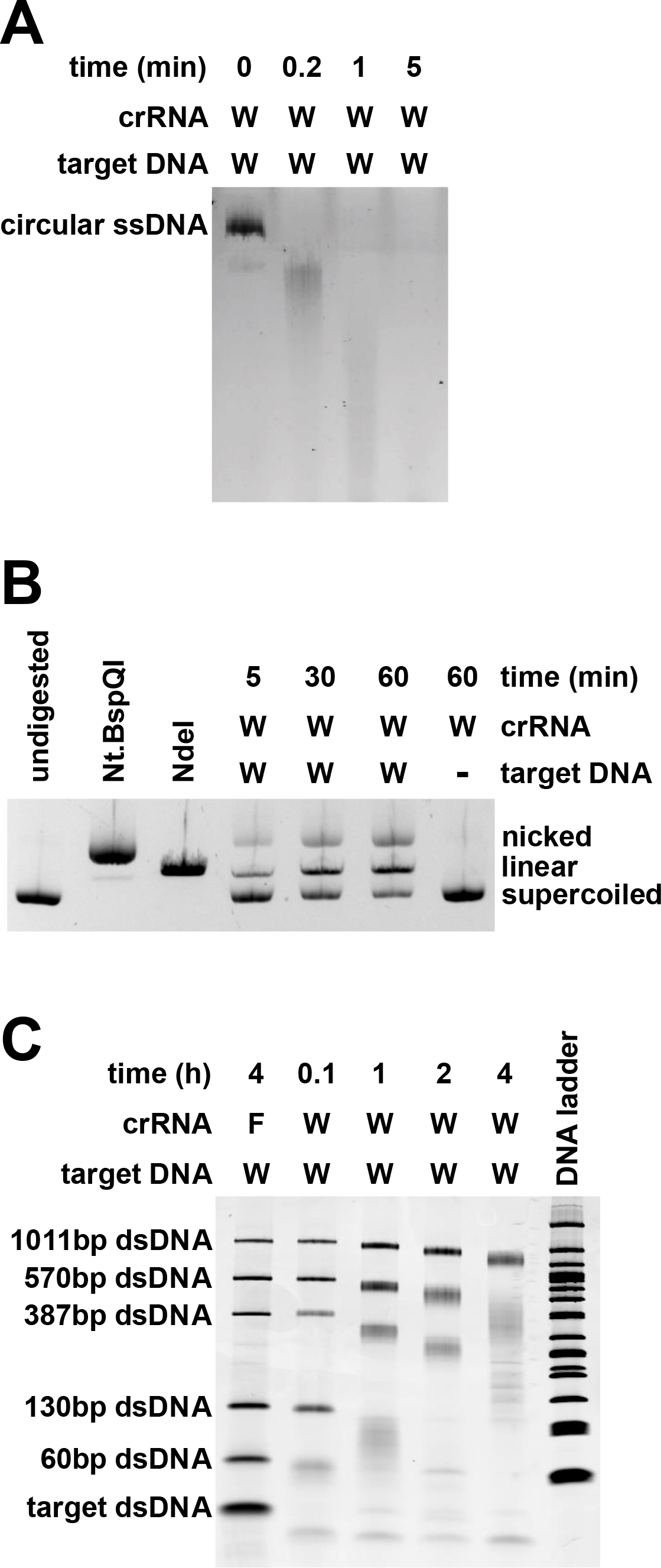
LbaCas12a *trans* nuclease activity on circular ssDNA, plasmid DNA, and linear dsDNA. LbaCas12a and a crRNA complementary to a short, dsDNA target were incubated in the presence of **(A)** circular M13mp18 ssDNA, **(B)** circular, supercoiled pUC19 dsDNA, or **(C)** five different non-targeted, linear dsDNAs of varying length. The pUC19 DNA was digested with restriction enzymes Nt.BspQI or NdeI (NEB) as controls for nicked or linear versions of the DNA, respectively. Sequences of crRNAs and substrates can be found in Supplementary Table S1. W = WTAP, F = FANCF.

**Figure 2.**
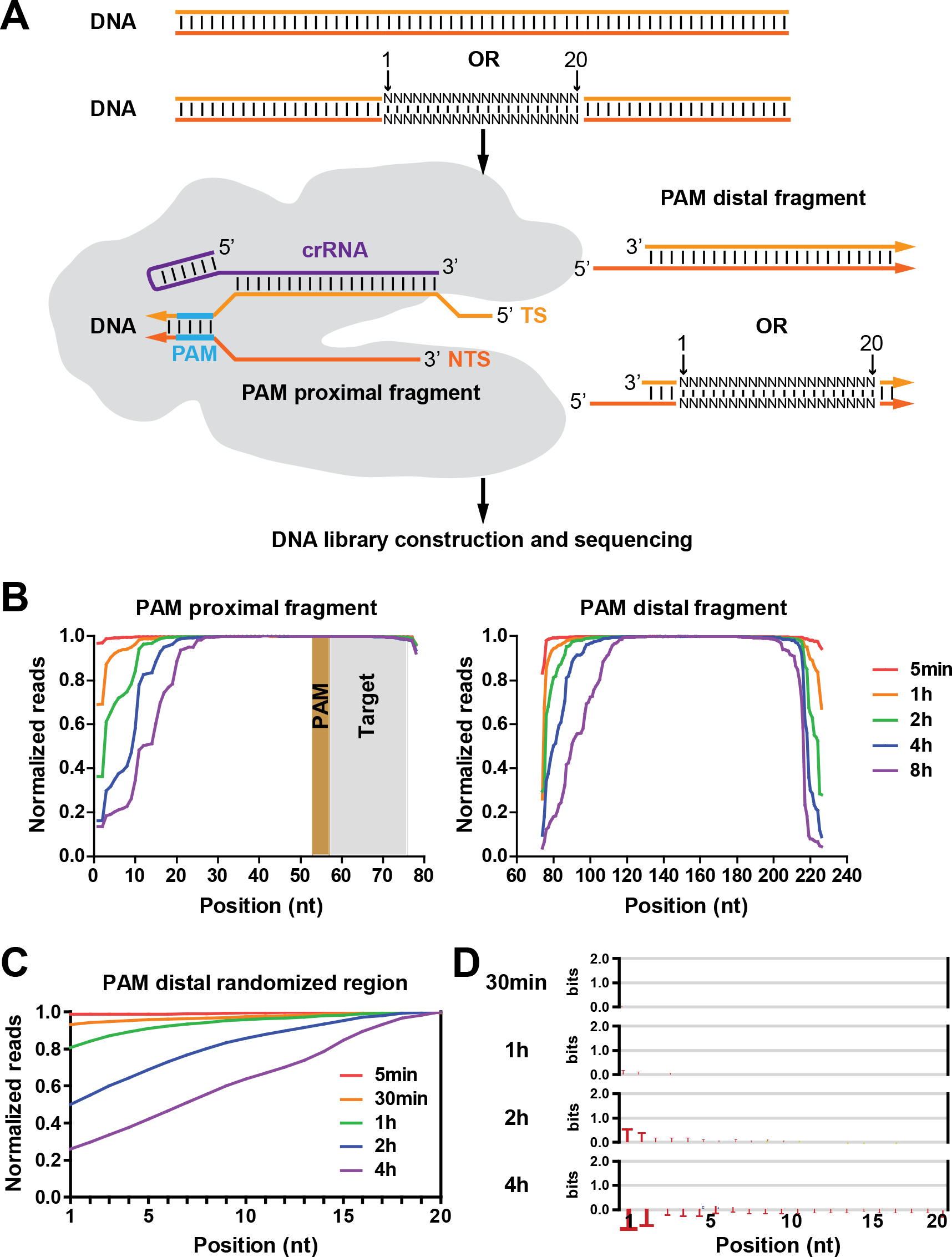
High-throughput sequencing characterization of LbaCas12a digestion reaction products. **(A)** Either a defined sequence dsDNA or a dsDNA containing a 20 nt long randomized region were subjected to digestion with LbaCas12a followed by DNA library construction and high-throughput sequencing. Aliquots of cleavage reactions were removed at indicated times, quenched and purified prior to DNA library construction. TS = target strand, NTS = non-target strand. **(B)** Normalized sequencing coverage of the PAM proximal and PAM distal pieces after cleavage of defined sequence dsDNA. The location of the PAM and target sequence within the PAM proximal piece of DNA are indicated by shaded areas. **(C)** Normalized sequencing coverage after cleavage of dsDNA containing a randomized region adjacent to the cleavage site. **(D)** Sequence logos showing preference for retention of specific bases at each position in the randomized region. A position weight matrix was made (see methods section) and plotted using the enoLOGOS web browser-based tool (32).

### Cas12a *trans* nuclease activity has minimal preference for primary sequence content of DNA

The observation that dsDNA was not immune to *trans* degradation via activated LbaCas12a led us to question what effect DNA sequence content might have on the degradation activity. Since ssDNA is a more susceptible substrate for the *trans* nuclease activity than dsDNA, we considered the possibility that regions of dsDNA with high A:T content transiently adopt single stranded states making them a better substrate for *trans* nuclease activity. In contrast, high G:C regions would be less prone to adopt a transient single-stranded state and thus be more resistant to cleavage. To test this possibility, we designed an *in vitro* cleavage assay using dsDNA produced by PCR that had a target sequence flanking a region of random sequence (Figure 2A). Care was taken to limit the PCR cycles so that the reaction was still in its exponential amplification phase (Supplementary Figure S1). This ensured that the substrate would be fully base paired rather than contain a mixture of mismatched indels. After cleavage of the target sequence, the PAM distal DNA fragment dissociates (24–26) and thus could become a substrate for *trans* cleavage. We took samples of a cleavage reaction at different time points and subjected the DNA that was present to Illumina sequencing. We observed an increasing loss of coverage in the random sequence region over time in a pattern consistent with loss at the DNA ends observed in the previous experiment (Figure 2C). However, the sequence content of the DNA remaining did not appreciably change (Figure 2D). The limited sequence content change is a small preference for the retention of T-rich sequences. We attribute this to be a PAM-dependent effect likely brought about by the protection from interrogating, guide-loaded but non-activated LbaCas12a molecules which are in excess. In addition, the slight accumulation of T’s is contradictory to the idea that lower stability A:T pairs are a better substrate for the *trans* activity. We interpret these results to indicate that Cas12a *trans* activity has no or very little preference for primary sequence content of non-target dsDNA.

### Target sequence identity and guide length influence *trans* nuclease activity

During our investigation we observed that not all activated LbaCas12a RNPs produced the same amount of *trans* activity. The guide and requisite target DNA pair used to activate the enzyme influenced how much *trans* activity was produced. An example of this is shown in Figure 3A, where the WTAP guide with 20 nt of target sequence and target DNA results in quicker nonspecific degradation of dsDNA than the FANCF guide with 20 nt of target sequence and target DNA. These results were produced despite the fact that the guide RNAs have a similar K_D_ when binding LbaCas12a (Figure 3B) and both have similar on-target activity which leads to the same amount of activated RNPs after 5 min (Supplemental Figure S2). We observed that guide RNA length also affected *trans* activity. Extending the targeting region of the guide RNA from 20 to 24 nt resulted in a noticeable reduction in *trans* activity on dsDNA and an apparent increase in activity on ssDNA (Figure 3C+3D). This is despite the fact that the guide RNA designs had a similar affinity for the Cas12a protein and produced similar on-target cleavage activity (Figure 3B, Supplementary Figure S2). The difference in *trans* activity is surprising because Cas12a enzymes are thought to interact with the first 20 nucleotides in the targeting portion of the guide while extensions past this region are thought to be unstructured/flexible and not involved in target recognition (25,33,34). We did not observe a noticeable difference in on-target cleavage activity when using a DNA substrate that either matched or did not match positions 21-24 in the guide RNA sequence, which is consistent with positions 21-24 not being involved in target recognition (Supplementary Figure S4).

**Figure 3.**
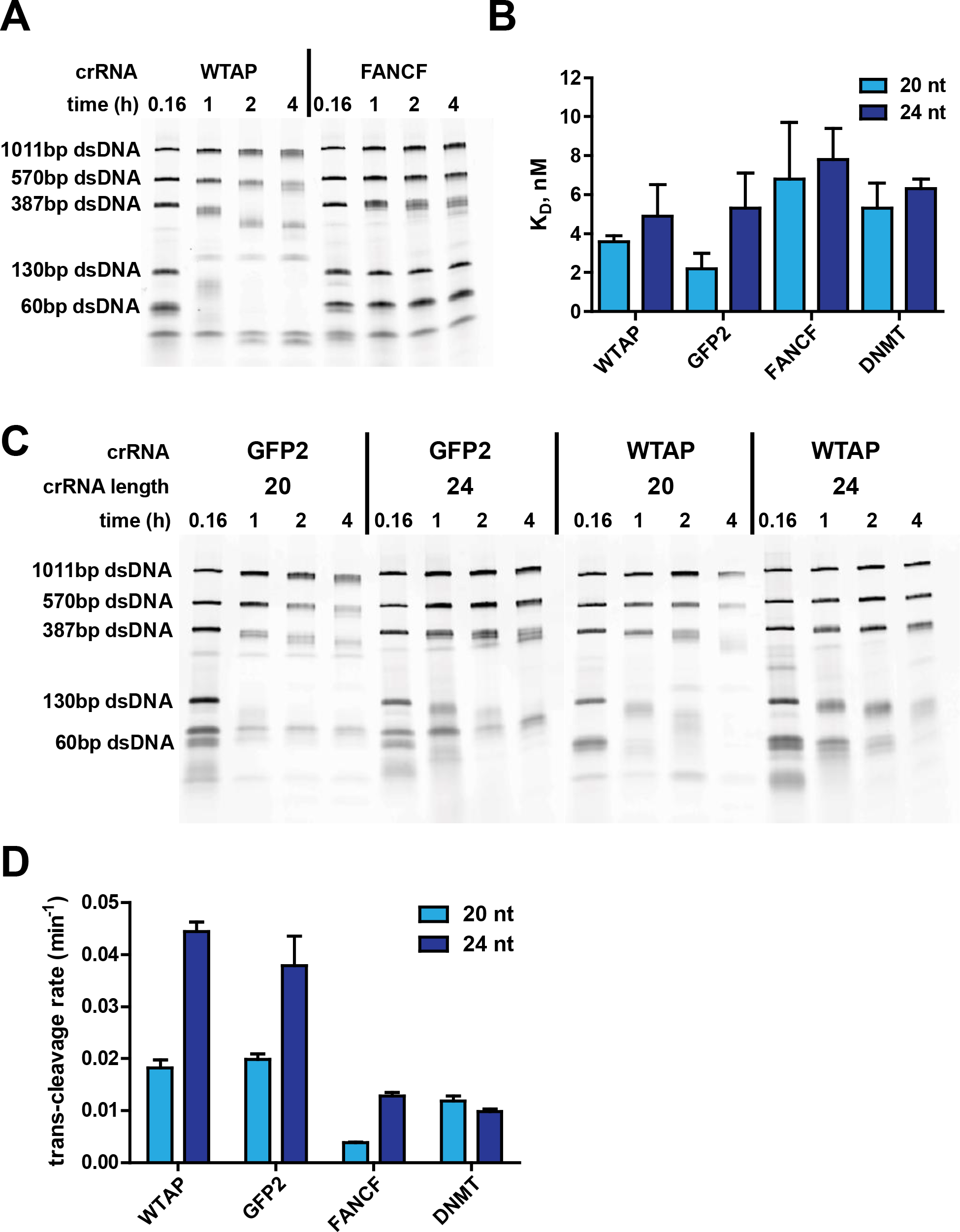
Guide RNA-dependent differences in *trans* nuclease activity. **(A)** Comparison of 2 different crRNAs with 20 nt target sequence length in a *trans* nuclease assay with dsDNA substrates. Each reaction contained a crRNA, LbaCas12a, on-target dsDNA and five non-target dsDNAs. W = WTAP, F = FANCF. **(B)** Dissociation constants of various crRNAs interacting with LbaCas12a. The crRNAs contain a targeting sequence that is either 20 or 24 nucleotides in length that targets one of 4 unique loci. **(C)** Comparison of crRNAs with a 20 versus 24 length targeting sequence in a *trans* nuclease assay with dsDNA substrates. Each reaction contained a crRNA, LbaCas12a, on-target dsDNA and five non-target dsDNAs. W = WTAP, G = GFP2. **(D)** Comparison of crRNAs with a 20 versus 24 length targeting sequence in a *trans* nuclease assay with a ssDNA substrate. Each reaction contained a crRNA, LbaCas12a, on-target dsDNA and a 5 nt long ssDNA with a fluorescein label on the 5’ end and an Iowa Black^®^ FQ quencher on the 3’ end. Raw traces of the data are provided in Supplementary Figure S3.

### Cas12a *trans* nuclease activity is influenced by magnesium and salt concentration

Since both on-target and *trans* activity depend upon the same RuvC domain, which uses Mg^2+^ in its cleavage mechanism (16,35,36), we investigated the effect of Mg^2+^ concentration in *in vitro* cleavage reactions. Reducing Mg^2+^ concentration noticeably slowed down degradation of a non-target dsDNA in *trans* (Figure 4A). At 2 mM Mg^2+^ there was little to no observable degradation of the DNA after 90 min. Although the on-target cleavage was likely slowed by the decreased Mg^2+^ as well, all of the on-target cleavage was still complete after 5 min even in 2 mM Mg^2+^. This suggests that reducing Mg^2+^ is an approach to minimize *trans* nuclease activity in a reactions of at least 5 min in length without sacrificing yield of on-target cleavage products. We also investigated the effect of monovalent salts on *trans* nuclease activity by titrating concentrations of NaCl and KCl in reactions with LbaCas12a. On-target DNA cleavage was slowed by increasing salt concentration, so care was taken to carry out reactions where on-target cleavage was complete for all samples to ensure an equal concentration of activated RNPs prior to addition of DNA for *trans* activity analysis (Supplementary Figure S5). The resulting *trans* degradation experiments show that the rate of dsDNA and ssDNA degradation noticeably slows as NaCl and KCl concentrations increase (Figures 4B,4C,5). We note that two studies describing *trans* nuclease activity utilized reactions where KCl concentrations were 150 mM or greater and pervasive degradation of off-target dsDNA was not observed (27,29). These results are consistent with our observations where *trans* nuclease activity on dsDNA is only clearly evident over reaction durations less than 1 hour when salt concentration is at or below 150 mM.

**Figure 4.**
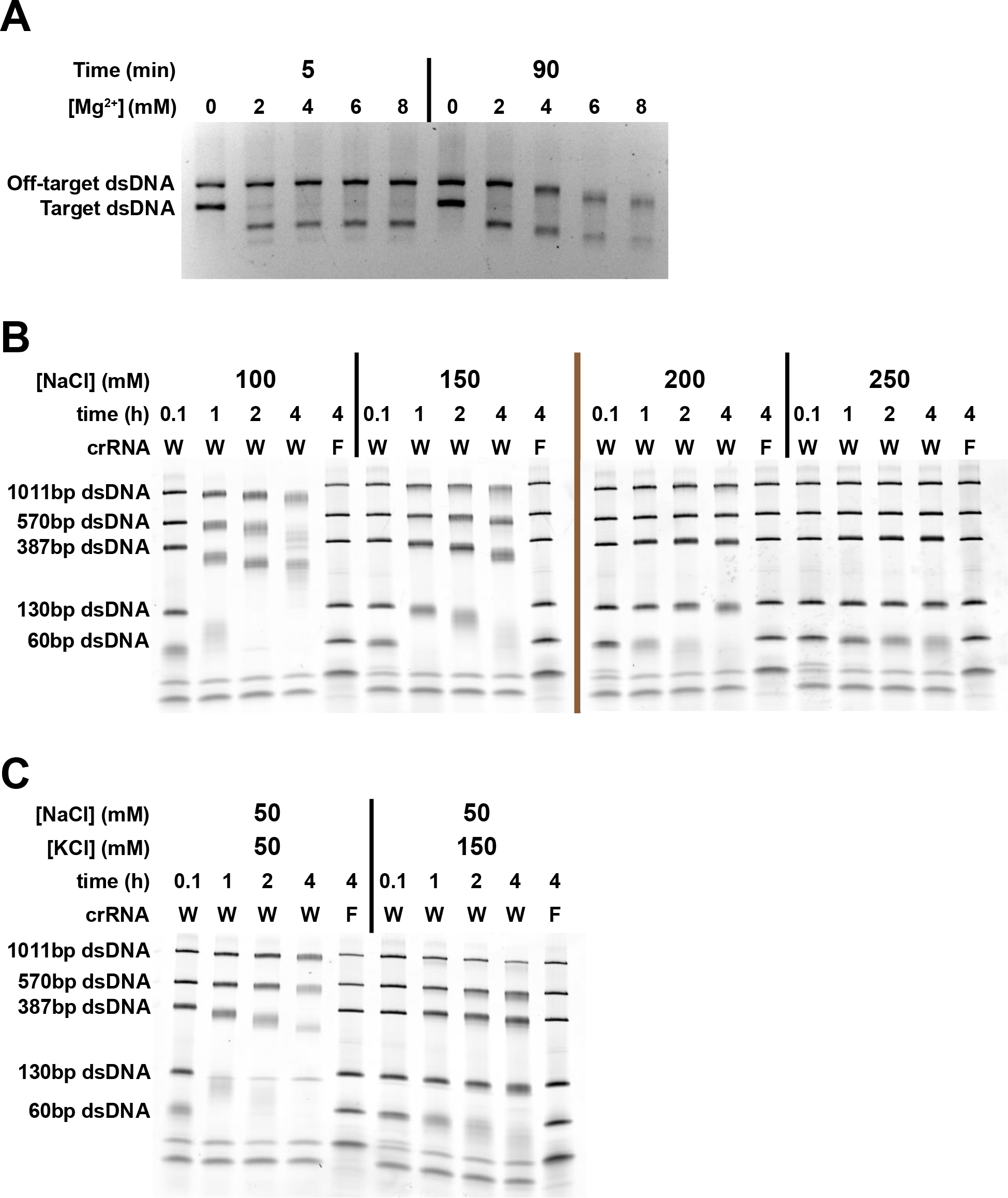
Reaction buffer composition modulates LbaCas12a activity. **(A)** Comparison of digestion reactions containing LbaCas12a, crRNA, an on-target and an off-target dsDNA with variable Mg^2+^ concentrations. **(B)** Comparison of digestion reactions with variable NaCl concentrations that contained LbaCas12a, crRNA, an on-target and five off-target dsDNAs. **(C)** Comparison of digestion reactions with variable NaCl and KCl concentrations that contained LbaCas12a, crRNA, an on-target and five off-target dsDNAs.

### RNA is also degraded by *trans* nuclease activity

In the course of our investigation, we suspected that RNA might be a substrate for the *trans* activity of Cas12a in addition to DNA. We conducted experiments where we substituted an RNA fluorescent reporter for its DNA counterpart. As shown in Figure 5, experiments with the 5 nt ssRNA reporter show that there is cleavage of the reporter, but only when LbaCas12a is activated by crRNA and matching target DNA. Notably, the rate of degradation for the ssRNA reporter was at least 10-fold slower than for ssDNA, confirming our observations that ssDNA is the most susceptible substrate for *trans* activity.

**Figure 5.**
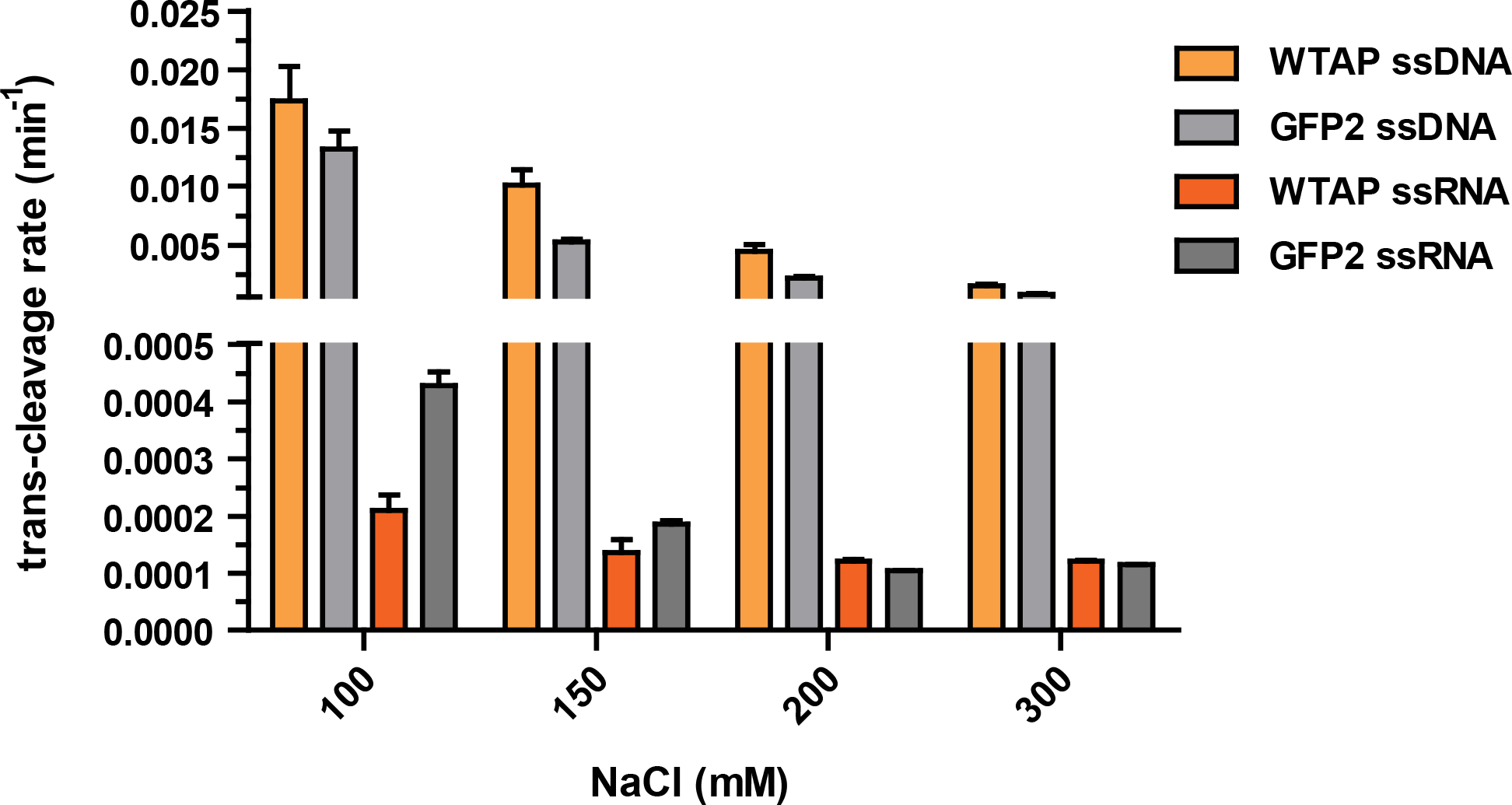
NaCl effect on *trans* nuclease activity with ssDNA and ssRNA substrates. Digestion reactions with variable NaCl concentrations that contained LbaCas12a, crRNA, an on-target dsDNA and a 5 nt long ssDNA or ssRNA with a fluorescein label on the 5’ end and an Iowa Black^®^ FQ quencher on the 3’ end were carried out and fluorescence was measured in 15 second intervals for either 1 hour (ssDNA) or 3 hours (ssRNA). The data were fit with a single exponential equation in Kaleidograph software to calculate the observed rate (see methods). Raw traces of the data are provided in Supplementary Figure S6.

### Lba, Asp and FnoCas12a orthologs have *trans* nuclease activity

Previous reports showed that all Cas12a orthologs tested thus far have *trans* nuclease activity (27,28). Consistent with previous findings, we observed *trans* nuclease activity with Lba, Asp, and FnoCas12a orthologs in cleavage assays using a fluorescent ssDNA reporter (Figure 6). Significantly, we observed differences in the level of *trans* nuclease activity between the orthologs when concentrations of activated RNPs were identical (Supplementary Figure S8). We consistently observed LbaCas12a to have the highest amount of *trans* activity of any Cas12a ortholog yet characterized. However, we cannot rule out the existence of a combination of reaction conditions (target sequence, buffer composition) where one of the other orthologs has enhanced activity.

**Figure 6.**
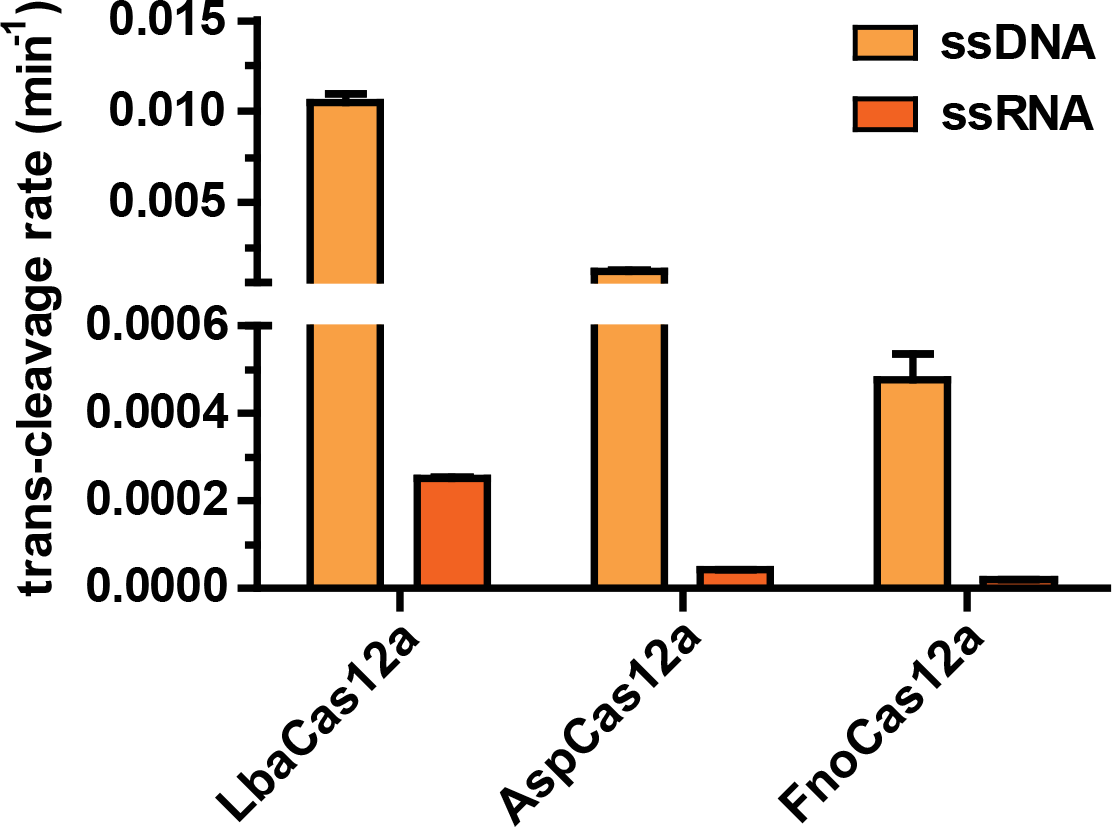
Comparison of the *trans* nuclease activity of Lba, Fno and AspCas12a. Digestion reactions contained crRNA, an on-target dsDNA, a 5 nt long ssDNA with a fluorescein label on the 5’ end and an Iowa Black^®^ FQ quencher on the 3’ end and either LbaCas12a, FnoCas12a or AspCas12a. Fluorescence was measured in 15 second intervals for either 2 hours (ssDNA) or 16 hours (ssRNA). The data were fit with a single exponential equation in Kaleidograph software to calculate the observed rate (see methods). Raw traces of the data are provided in Supplementary Figure S7.

## DISCUSSION

Like Cas9 proteins from Class 2 Type II CRISPR systems, Cas12a proteins from Class 2 Type V-A CRISPR systems have been used for genome editing in cells and DNA cleavage applications *in vitro*. Both Cas9 and Cas12a carry out DNA cleavage via a single-turnover, RNA-guided mechanism. However, extensive differences exist in the details of how the two enzyme types carry out this function. This list of differences was expanded with the recent observation that target DNA-bound Cas12a RNP remains in an activated state post cleavage. This unleashes multiple turnover, non-specific degradation of ssDNA in *trans* via the RuvC domain (27–29). Structural studies have noted that ssDNA should be the only substrate for *trans* nuclease activity as it can fit in the active site of the FnoCas12a RNP complex, but dsDNA would have steric clashes (29). Despite this, we observe that LbaCas12a is able to degrade dsDNA, in *trans*, at a slower rate than ssDNA. Although the rate for dsDNA degradation is much slower, there is the potential for significant damage of dsDNA in *trans* over the course of typical *in vitro* reactions. This is especially true if the reaction conditions consist of monovalent salt concentration of <150 mM and Mg^2+^ concentrations >2 mM, conditions under which we observe LbaCas12a to be most active. Other studies involving *trans* nuclease activity have utilized reaction conditions where salt concentrations were ≥150 mM, making it difficult to observe dsDNA degradation over the duration of typical reactions.

Our results show that Cas12a *trans* nuclease activity can nick circular double-stranded plasmid DNA. We interpret this to mean that *trans* nuclease activity likely is nicking dsDNA rather than making discreet double-stranded breaks. This is consistent with the model of a transient interaction of a non-target DNA in *trans* with the activated RuvC domain. To carry out a double stranded break of the non-target DNA, structural rearrangements would be necessary to cleave both strands of the transiently interacting DNA, an unlikely possibility given that Cas12a remains bound to its guide RNA and target DNA during the process. Pervasive nicking by multiple turnover events on dsDNA would result in the DNA falling apart as sites of nicking accumulate across from each other or close to the ends of linear DNA.

It was proposed that the small amount of dsDNA degradation observed in previous studies was due to breathing at the ends of linear dsDNA causing single strands to be transiently available for the nuclease to act on. While we cannot rule end-breathing out as being a factor entirely, we observed no preference for the retention of high G:C regions at DNA ends in a *trans* cleavage reaction. We interpret this to mean that DNA breathing is not a major contributor to the susceptibility of dsDNA to the *trans* nuclease activity of Cas12a orthologs.

Salt concentration differences can alter the stability of annealed DNA strands. However, within the range of Mg^2+^ and Na^+^ or K^+^ concentration used in this study the Mg^2+^ concentration is the driving determinant of DNA melting temperature (37). The effects of Na^+^ or K^+^ concentration on DNA duplex stability are negligible. It is likely that the salt effects on *trans* nuclease activity that we observed were due to modulation of the electrostatic potential required for nucleic acids to interact with the activated RNP in *trans*, i.e. in lower salt concentrations transient electrostatic interactions are more favorable than in higher salt concentrations.

Our data show that in addition to ssDNA and dsDNA, RNA is also a substrate for Cas12a *trans* nuclease activity. The rate of RNA degradation was at least 20-fold slower than for ssDNA in all cases.

Taken together, our findings allow us to update the model of Cas12a *trans* nuclease activity to show that although they vary in susceptibility based on the factors described above, no nucleic acids are completely safe from *trans* nuclease activity (Figure 7).

**Figure 7.**
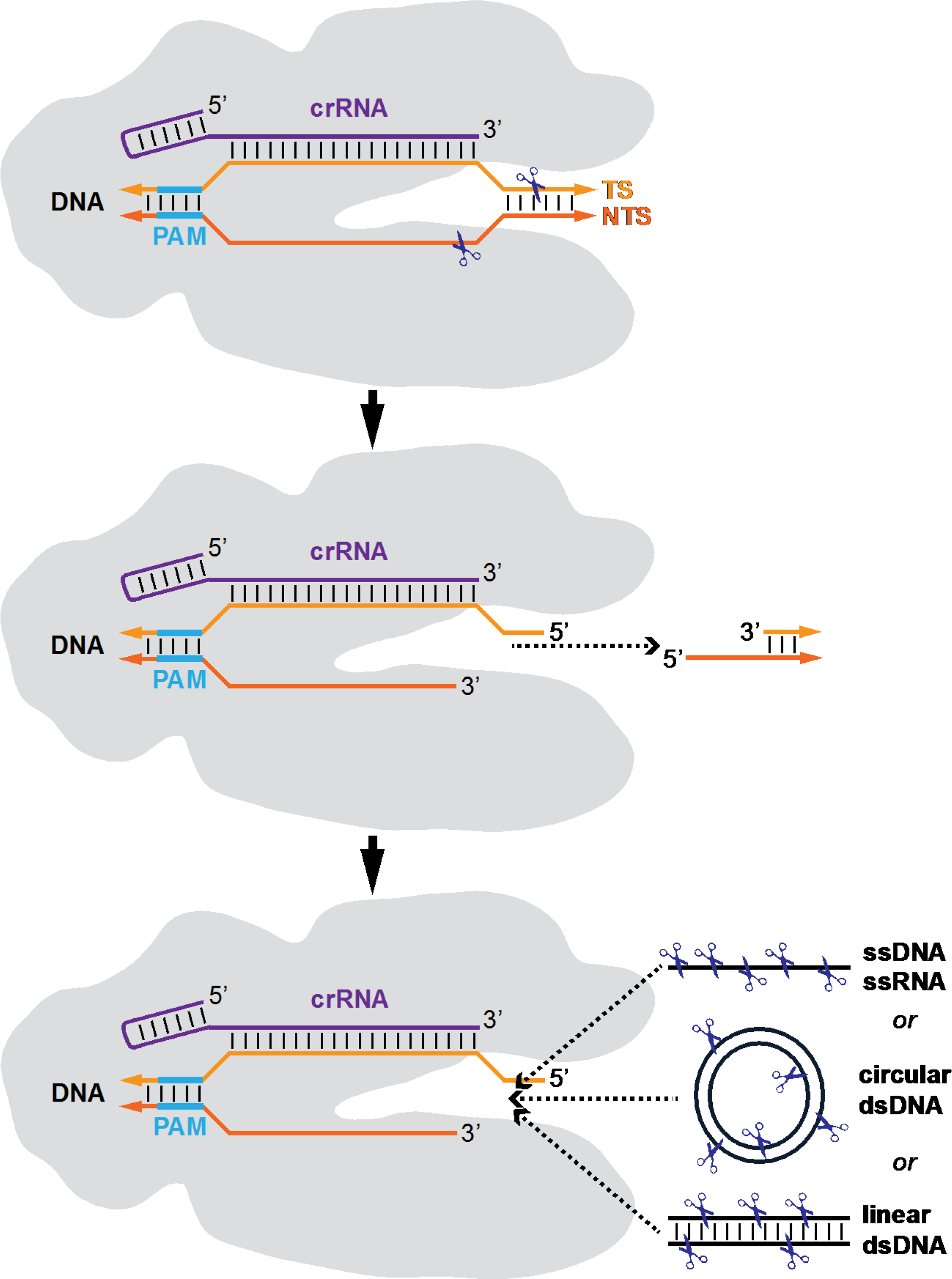
An updated model for Cas12a activity. Cas12a-crRNA RNP binds to a target DNA that is complementary to the crRNA targeting sequence. Both strands of the target DNA are cleaved and the PAM distal piece of DNA dissociates. The PAM proximal piece remains bound and this results in the RNP remaining in an activated state and nucleic acids, with ssDNA being the most susceptible, can transiently interact with the active site and be cleaved or nicked.

For particular *in vitro* applications, such as cleavage to make compatible 5’ overhangs for cloning or cleavage of a ssDNA reporter in *trans* to detect low abundance targets (27,38) it would be desirable to have the *trans* activity of the Cas12a enzyme to be as low as possible or as high as possible, respectively. We observe that for LbaCas12a, the on-target reaction is essentially complete after 5 min even in the presence of Mg^2+^ concentrations as low as 2 mM. Although reaction conditions may need to be adjusted for given guide-target combinations and different Cas12a orthologs, in general keeping reaction times short and Mg^2+^ concentration low will reduce the impact of *trans* activity while the opposite will increase it. Our results also show that the activity can be modulated by guide RNA design choices. The FANCF guide/target combination and WTAP guide/target combination that were used in this study produced very different levels of *trans* activity despite having the similar guide RNA-LbaCas12a binding affinity and similar on-target cleavage activity. This indicates that another way to modulate activity resides within the nature of the target sequence itself. The explanation for our observation remains elusive and difficult to test in a logical, high-throughput manner and will require future investigation.

Guide RNA spacer (target) lengths of 20 nt tend to produce higher *trans* activity than RNAs that are extended by 4 more nt and the extended sequence does not have to match the target DNA for this effect. Notably, published structural data have shown that Cas12a orthologs do not utilize the guide RNA for target recognition after base 20 (25,33,34). It is likely that the longer 3’ end sterically inhibits *trans* DNA interacting with the RuvC domain. Future studies will be required to determine the exact mechanism. It would be interesting to learn what effect recently proposed Cas12a guide RNA design changes and modifications (39–42) have on *trans* activity. In addition to guide RNA design, we find that the choice of Cas12a ortholog to use in a reaction will greatly influence how much *trans* activity is present. LbaCas12a appears to be the ortholog of choice for high *trans* activity applications. However, recent studies have expanded the known diversity and classification of Cas12 systems and have introduced mutant Cas12 proteins with altered PAM specificities (43–47). There is the possibility that there may be other orthologs with very high or very low *trans* activity and future studies will be needed to determine why orthologs vary in magnitude of *trans* activity.

An important question remaining regarding *trans* nuclease activity is what effect it has on genome editing in cells. Although LbaCas12a produces high *trans* activity, it works well in genome editing in mammalian cells and is reported as producing a very low level of off-target edits (48,49). Additionally, LbaCas12a is the Cas12a of choice for genome editing in organisms that are grown at temperatures below 30°C because AspCas12a has minimal activity below 30°C (50). Together, these successes indicate that *trans* nuclease activity does not have catastrophic or even noticeable effects on the genomic stability of cells from a variety of organisms. Potential explanations might include: removal of the RNP from the target DNA post-cleavage by cellular repair machinery before *trans* activity causes pervasive damage, protection of cellular DNA by histones and other proteins, and mutation-free repair of DNA nicks. Ultimately, it will be difficult to isolate and study the actual effect of *trans* nuclease activity on genome editing since it is reliant on the same RuvC active site as the on-target activity. Perhaps the careful timing of introducing inhibitors to turn off or reset the Cas12a RNP will can be used to distinguish the effects of on-target and *trans* activity in cells (51,52). It will be important to support future work so that the ramifications of Cas12a *trans* nuclease activity in genome editing applications can be fully understood.

## ACKNOWLEDGEMENTS

We thank Laurie Mazzola, Darcie Spaulding, Danielle Fuchs and Aine Quimby for assistance with Illumina sequencing and capillary electrophoresis.

