## SupplementaryMaterials for "Cas12a trans-cleavage can be modulated in vitro and is active on ssDNA, dsDNA, and RNA"

Supplementary Material

Supplementary Table S1. DNA and RNA sequences

| Name | Type | Source | Sequence | Notes |
|---|---|---|---|---|
| AspCas12a WTAP 20 crRNA | RNA | IDT | UAAUUUCUACUCUUGUAGAUCCACUCACUG CUUUCUCCUC | |
| FnoCas12a WTAP 20 crRNA | RNA | IDT | UAAUUUCUACUGUUGUAGAUCCACUCACUG CUUUCUCCUC | |
| LbaCas12a WTAP 20 crRNA | RNA | IDT | UAAUUUCUACUAAGUGUAGAUCCACUCACU GCUUUCUCCUC | |
| LbaCas12a WTAP 24 crRNA | RNA | IDT | UAAUUUCUACUAAGUGUAGAUCCACUCACU GCUUUCUCCUCUUGA | |
| LbaCas12a DNMT 20 crRNA | RNA | IDT | UAAUUUCUACUAAGUGUAGAUUUUCCCUUC AGCUAAAAUAA | |
| LbaCas12a DNMT 24 crRNA | RNA | IDT | UAAUUUCUACUAAGUGUAGAUUUUCCCUUC AGCUAAAAUAAAGGA | |
| LbaCas12a FANCF 20 crRNA | RNA | IDT | UAAUUUCUACUAAGUGUAGAUGUCGGCAUG GCCCCAUUCGC | |
| LbaCas12a FANCF 24 crRNA | RNA | IDT | UAAUUUCUACUAAGUGUAGAUGUCGGCAUG GCCCCAUUCGCACGG | |
| LbaCas12a GFP2 20 crRNA | RNA | Sigma | UAAUUUCUACUAAGUGUAGAUACAAGCAGA AGAACGGCAUC | |
| LbaCas12a GFP2 24 crRNA | RNA | Sigma | UAAUUUCUACUAAGUGUAGAUACAAGCAGA AGAACGGCAUCAAGG | |
| DNMT randomized template | DNA | IDT | CTGTTACTCGCCTGTCAAGTGGCGTGACAC CGGGCGTGTTCCCCAGAGTGACTTTTCCTT TTATTTCCCTTCAGCTAAAATAAAGGANNN NNNNNNNNNNNNNNNNNAGTTCTGCCCTCC CGTCACCCCTGTTTCTGGCACCAGGAATCC CCAACATGCACTGATGTTGTGTTTTTAACA TGTCAATCTGTCCGTTCACA | *Used in PCR to make substrate for Figure 2C+2D |
| DNMT Fwd primer | DNA | IDT | CTGTTACTCGCCTGTCAAGTGG | *Used in PCR to make substrate for Figure 2C+2D |
| DNMT Rev primer | DNA | IDT | TGTGAACGGACAGATTGACATGT | *Used in PCR to make substrate for Figure 2C+2D |
| WTAP small activator Top | DNA | IDT | ATTACCTTTCCCACTCACTGCTTTCTCCTC TTGAG | *Anneal to make small activator DNAs |
| WTAP small activator Bottom | DNA | IDT | CTCAAGAGGAGAAAGCAGTGAGTGGGAAAG GTAAT | *Anneal to make small activator DNAs |
| DNMT small activator Top | DNA | IDT | TTTCCTTTTATTTCCCTTCAGCTAAAATAA AGGAG | *Anneal to make small activator DNAs |
| DNMT small activator Bottom | DNA | IDT | CTCCTTTATTTTAGCTGAAGGGAAATAAAA GGAAA | *Anneal to make small activator DNAs |
| FANCF small activator Top | DNA | IDT | CGGCGCTTTGGTCGGCATGGCCCCATTCGC ACGGC | *Anneal to make small activator DNAs |
| FANCF small activator Bottom | DNA | IDT | GCCGTGCGAATGGGGCCATGCCGACCAAAG CGCCG | *Anneal to make small activator DNAs |

| | | | | |
|---|---|---|---|---|
| GFP2 small activator Top | DNA | IDT | CATGCATTTGACAAGCAGAAGAACGGCATCAAGGG | *Anneal to make small activator DNAs |
| GFP2 small activator Bottom | DNA | IDT | CCCTTGATGCCGTTCTTCTGCTTGTCAAATGCATG | *Anneal to make small activator DNAs |
| WTAP activator template 20 | DNA | IDT | ACCTGCCAACCAAAGCGAGAACCGCTATTGCTAGCCATTTACCACTCACTGCTTTCTCCTCTCCGATCCTATATTACGCTCATGAGACAATAACCCTGA | |
| WTAP activator template 24 | DNA | IDT | ACCTGCCAACCAAAGCGAGAACCGCTATTGCTAGCCATTTACCACTCACTGCTTTCTCCTCTTGATCCGATCCTATATTACGCTCATGAGACAATAACCCTGA | |
| FANCF activator template 20 | DNA | IDT | ACCTGCCAACCAAAGCGAGAACCGCTATTGCTAGCCATTTAGTCGGCATGGCCCCATTCGCTCCGATCCTATATTACGCTCATGAGACAATAACCCTGA | |
| FANCF activator template 24 | DNA | IDT | ACCTGCCAACCAAAGCGAGAACCGCTATTGCTAGCCATTTAGTCGGCATGGCCCCATTCGCACGGTCCGATCCTATATTACGCTCATGAGACAATAACCCTGA | |
| DNMT activator template 20 | DNA | IDT | ACCTGCCAACCAAAGCGAGAACCGCTATTGCTAGCCATTTATTTCCCTTCAGCTAAAATAATCCGATCCTATATTACGCTCATGAGACAATAACCCTGA | |
| DNMT activator template 24 | DNA | IDT | ACCTGCCAACCAAAGCGAGAACCGCTATTGCTAGCCATTTATTTCCCTTCAGCTAAAATAAAGGATCCGATCCTATATTACGCTCATGAGACAATAACCCTGA | |
| GFP2 activator template 24 | DNA | IDT | ACCTGCCAACCAAAGCGAGAACCGCTATTGCTAGCCATTTAACAAGCAGAAGAACGGCATCAAGGTCCGATCCTATATTACGCTCATGAGACAATAACCCTGA | |
| Activator template Fwd primer | DNA | IDT | ACCTGCCAACCAAAGCGAGAAC | *PCR from templates to make activator DNAs |
| Activator template Rev primer | DNA | IDT | TCAGGGTTATTGTCTCATGAGCG | *PCR from templates to make activator DNAs |
| DNA fluorescent reporter | DNA | IDT | /56-FAM/NNNNN/3IABkFQ/ | |
| RNA fluorescent reporter | RNA | IDT | /56-FAM/NNNNN/3IABkFQ/ | |
| WTAP amplicon Fwd primer | DNA | IDT | TCCAACAGCTCAGAGGAGAGAAC | *PCR from HEK-293 DNA to make substrate for Figure 2B |
| WTAP amplicon Rev primer | DNA | IDT | CACTTGAGTCCAAGCCATTCTG | *PCR from HEK-293 DNA to make substrate for Figure 2B |
| WTAP 226bp amplicon | DNA | PCR | TCCAACAGCTCAGAGGAGAGAACTGGCAGAGGAGGTAGTGGTTACGTAAATCAACTCAGTGCGGGGTATGAAAGTGTAGACTCTCCCACGGGCAGTGAAAACTCTCTCACACACCAATCAAATGACACAGACTCCAGTCATGACCCTCAAGAGGAGAAAGCAGTGAGTGGGAAAGGTAATCGAACTGTGGGTTCCCGCCACGTTCAGAATGGCTTGGACTCAAGTG | Top strand sequence of PCR product made by Fwd and Rev primers using HEK-293 gDNA as template. PCR product was used as a substrate for Figure 2B |

| | | | | |
|---|---|---|---|---|
| DNA ladder 60bp | DNA | IDT | CCTTGCATCGTTCGACTGGGTTTGTTC<br>TACATAAAACGGGCGCAACGTCTGCTC<br>TGGTCA | Top strand sequence of annealed oligos used as off-target dsDNA in Figures 1C,2A,2C,4B,4C |
| DNA ladder 130bp | DNA | PCR | CACGACAGGATAGACGACAACGCCCAC<br>AGCCACCTGAGGGCGAGCCTGCTCGGT<br>GCGAGCGAGTGCTTCCCAGTGGTGGAT<br>GGAAGGCTGGTAAGGGGGACGTGGCAG<br>CAGATATTCTTCGTCGAGCTCG | Top strand sequence of PCR product used as off-target dsDNA in Figures 1C,2A,2C,4B,4C |
| DNA ladder 387bp | DNA | PCR | ACCTGCCAACCAAAGCGAGAACATGGG<br>AGCAGCTGGTCAGAGGGGACCCCGGCC<br>TGGGGCCCCTAACCCTATGTAGCCTCA<br>GTCTTCCCATCAGGCTCTCAGCTCAGC<br>CTGAGTGTTGAGGCCCCAGTGGCTGCT<br>CTGGGGGCCTCCTGAGTTTCTCATCTG<br>TGCCCCTCCCTCCCTGGCCCAGGTGAA<br>GGTGTGGTTCCAGAACCGGAGGACAAA<br>GTACAAACGGCAGAAGCTGGAGGAGGA<br>AGGGCCTGAGTCCGAGCAGAAGAAGAA<br>GGGCTCCCATCACATCAACCGGTGGCG<br>CATTGCCACGAAGCAGGCCAATGGGGA<br>GGACATCGATGTCACCTCCAATGACTA<br>GGGTGGGCAACCACGCTCATGAGACAA<br>TAACCCTGA | Top strand sequence of PCR product used as off-target dsDNA in Figures 1C,2A,2C,4B,4C |
| DNA ladder 570bp | DNA | PCR | CCGCATGCGCGTCTATAAGCAACATGG<br>CCGCTCTGCCAGTCACCAACATGGCAA<br>CCACAGCCTCACTGTGTCCGGAAGCAT<br>GAGAGAGCGGGTATGAGGACTATACCG<br>GCCGCTCTGCCAACACCTAACGTGGCT<br>CTTTATAGCTTCCAAGAAAGCGCGACA<br>CCGGAAATGACTTTTCTGTCTTGCTCA<br>GCTCCAGGGGTCATTTTCCGGTTAGCC<br>TTCGGGGTGTCCGCGTGAGAATTGGCT<br>ATATCCTGGAGCGAGTGCTGGGAGGTG<br>CTAGTCCGCCGCGCCTTATTCGAGAGG<br>TGTCAGGGCTGGGAGACTAGGATGTCG<br>GACACGTGGAGCTCTATCCAGGCCCAC<br>AAGAAGCAGCTGGACTCTCTGCGGGAG<br>AGGCTGCAGCGGAGGCGGAAGCAGGAC<br>TCGGGGCACTTGGGTGAGGCACTGGGC<br>TGTTGGGGCCGAGGCTCAACCGGGGAG<br>GACGGGCGGGATCCCTAGAGAAAAAGC<br>TCCAGAGTCACTGCCCTTTACCAAACG<br>GCCATTTCTATAAAGCCTCGTACATTG<br>GTACTCCCTCTCCACCTTCAGCAGCAG<br>ACG | Top strand sequence of PCR product used as off-target dsDNA in Figures 1C,2A,2C,4B,4C |
| DNA ladder 1011bp | DNA | PCR | CATTCAGGCTGCGCAACTGTTGGGAAG<br>GGCGATCGGTGCGGGCCTCTTCGCTAT<br>TACGCCAGCTGGCGAAAGGGGGATGTG<br>CTGCAAGGCGATTAAGTTGGGTAACGC<br>CAGGGTTTTCCCAGTCACGACGTTGTA | Top strand sequence of PCR product used as off-target dsDNA in Figures 1C,2A,2C,4B,4C |

| | | | |
|---|---|---|---|
| | | | AAACGACGGCCAGTGAATTCGAGCTCG<br>GTACCCGGGGATCCTCTAGAGTCGACC<br>TGCAGGCATGCAAGCTTGGCGTAATCA<br>TGGTCATAGCTGTTTCCTGTGTGAAAT<br>TGTTATCCGCTCACAATTCCACACAAC<br>ATACGAGCCGGAAGCATAAAGTGTAAA<br>GCCTGGGGTGCCTAATGAGTGAGCTAA<br>CTCACATTAATTGCGTTGCGCTCACTG<br>CCCGCTTTCCAGTCGGGAAACCTGTCG<br>TGCCAGCTGCATTAATGAATCGGCCAA<br>CGCGCGGGGAGAGGCGGTTTGCGTATT<br>GGGCGCTCTTCCGCTTCCTCGCTCACT<br>GACTCGCTGCGCTCGGTCGTTCGGCTG<br>CGGCGAGCGGTATCAGCTCACTCAAAG<br>GCGGTAATACGGTTATCCACAGAATCA<br>GGGGATAACGCAGGAAAGAACATGTGA<br>GCAAAAGGCCAGCAAAAGGCCAGGAAC<br>CGTAAAAAGGCCGCGTTGCTGGCGTTT<br>TTCCATAGGCTCCGCCCCCCTGACGAG<br>CATCACAAAAATCGACGCTCAAGTCAG<br>AGGTGGCGAAACCCGACAGGACTATAA<br>AGATACCAGGCGTTTCCCCCTGGAAGC<br>TCCCTCGTGCGCTCTCCTGTTCCGACC<br>CTGCCGCTTACCGGATACCTGTCCGCC<br>TTTCTCCCTTCGGGAAGCGTGGCGCTT<br>TCTCATAGCTCACGCTGTAGGTATCTC<br>AGTTCGGTGTAGGTCGTTCGCTCCAAG<br>CTGGGCTGTGTGCACGAACCCCCCGTT<br>CAGCCCGACCGCTGCGCCTTATCCGGT<br>AACTATCGTCTTGAGTCCAACCCGGTA<br>AGACACGACTTATCGCCACTGGCAGCA<br>GCCACTGGTAACAGGATTAGCAGAGCG<br>AGGTATGTAGGC |

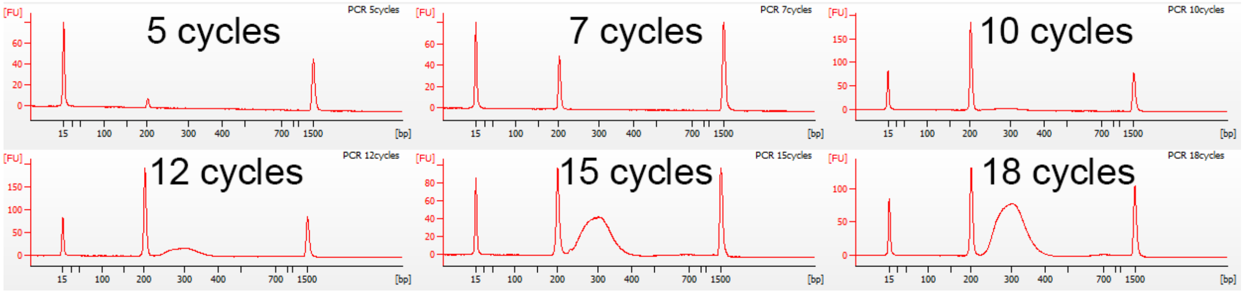

Supplementary Figure S1. PCR cycle titration using DNMT randomized template DNA. A PCR reaction was set up in 1X Q5® Hot Start High-Fidelity 2X Master Mix (NEB #M0494) with 0.055 pmol of DNMT randomized template, 50 pmol each of DNMT Fwd primer and DNMT Rev primer in a total volume of 100 µl. Reactions were incubated for 30 seconds at 98°C and then cycled either 5, 7, 10, 12, 15, or 18 times at the following temperatures: 98°C 10 sec, 65°C 15 sec, 72°C 15 sec. PCR reactions were purified by column purification, loaded on a DNA1000 chip, and run on an Agilent 2100 Bioanalyzer instrument. Expected product size was 200 bp, a smear of indels centered ~300 bp becomes prominent at 12 cycles and greater. PCR product after 7 cycles of amplification was chosen for use as a substrate for the experiments in Figure 2.

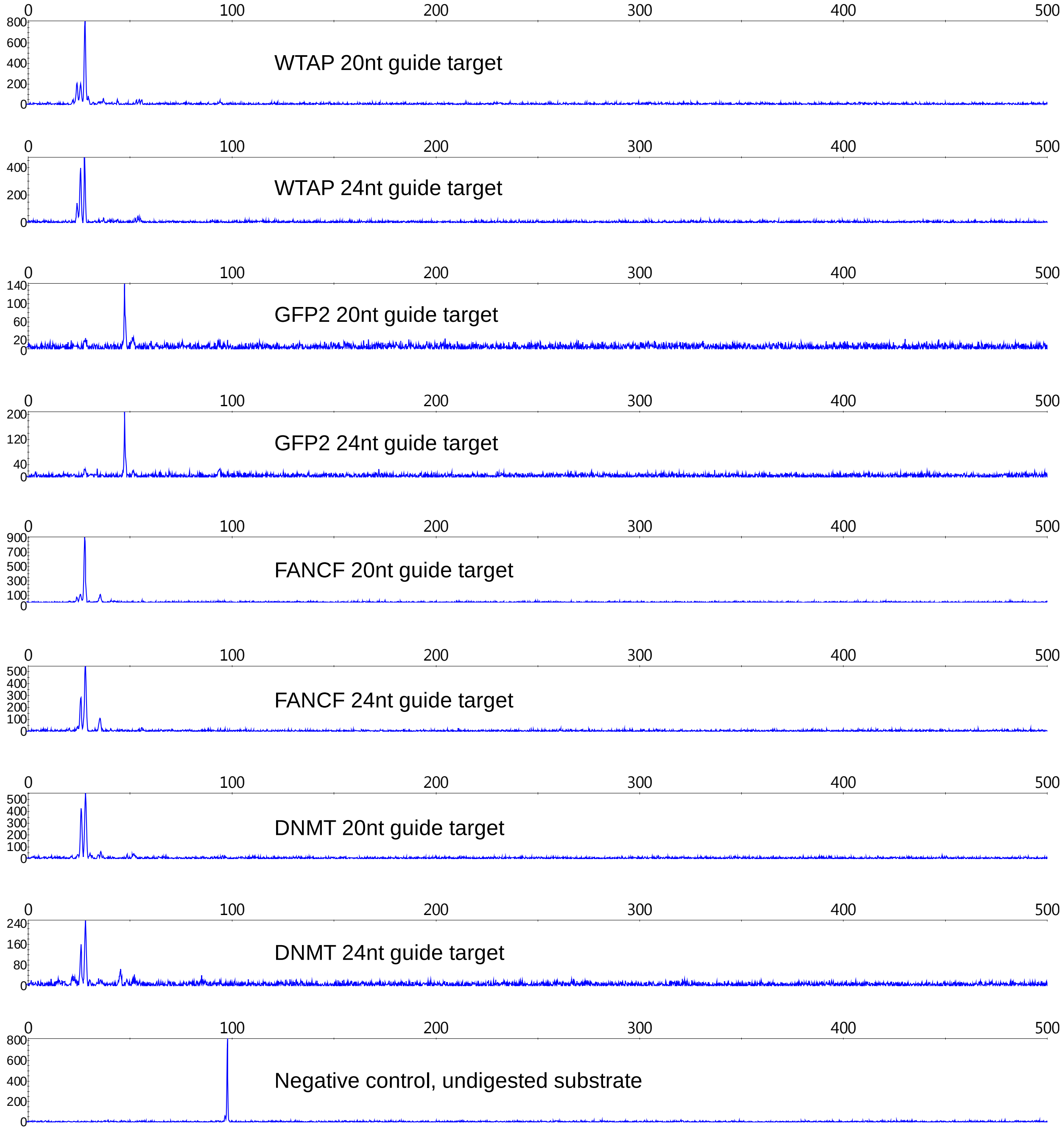

Supplementary Figure S2. Cleavage of on-target DNA substrates by LbaCas12a with various guide RNA and target combinations. RNPs were formed in NEB Buffer 2.1 for 10 min at room temperature. Substrate DNA with a 5' FAM label was added and samples were incubated for 5 min at room temperature. Reactions were quenched by adding EDTA and samples were column purified before being analyzed by capillary electrophoresis.

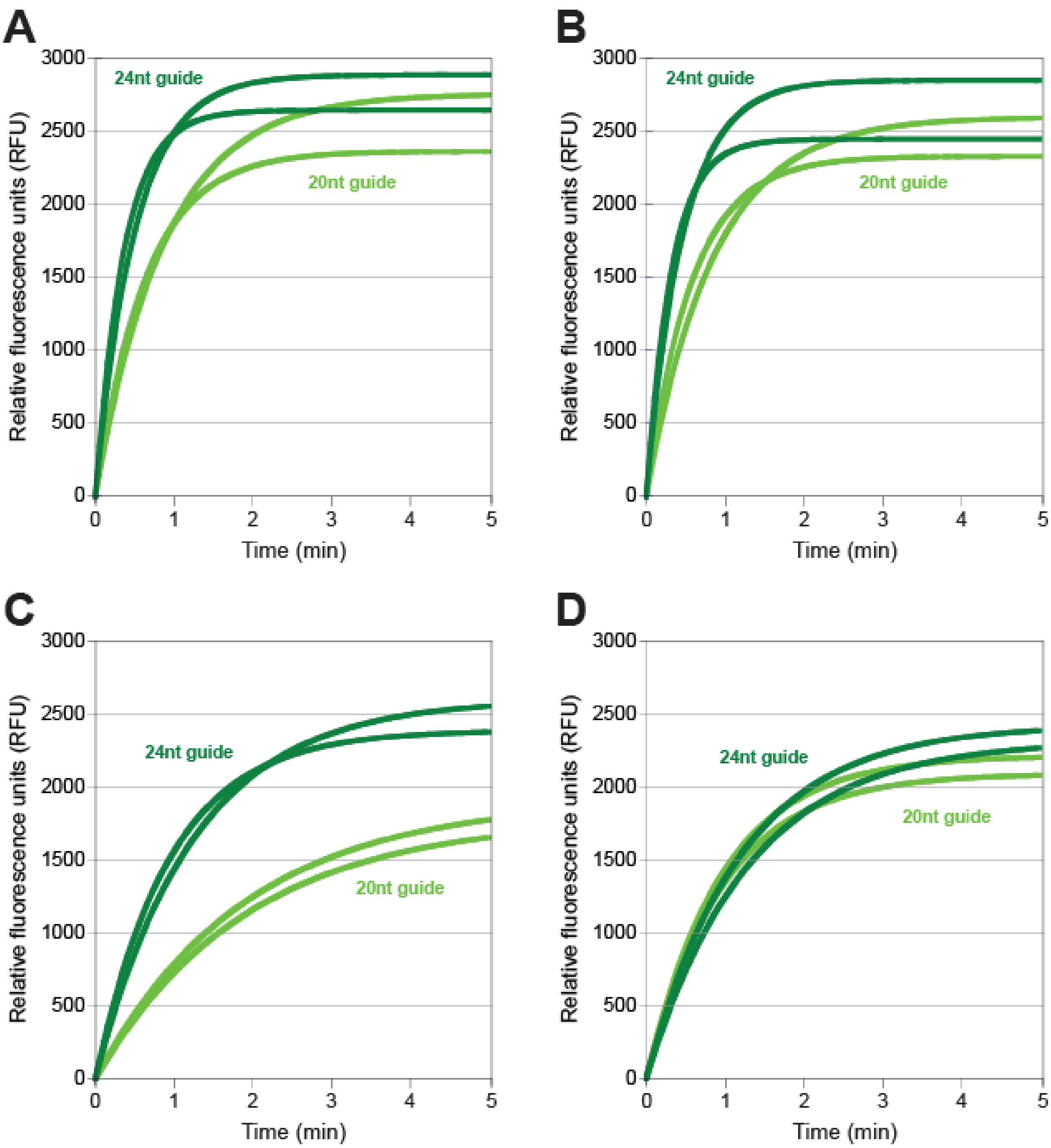

Supplementary Figure S3. Examples of raw traces from fluorescent ssDNA reporter assay. **(A)** WTAP guides and activator DNA, **(B)** GFP2 guides and activator DNA, **(C)** FANCF guides and activator DNA, **(D)** DNMT guides and activator DNA.

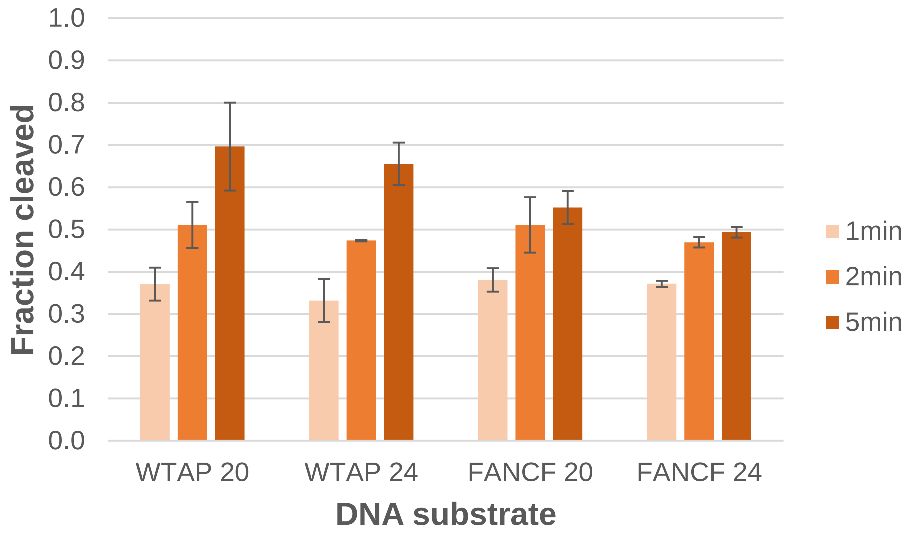

Supplementary Figure S4. Cleavage of DNA substrates with either 20 or 24 bases that match a guide RNA targeting region. RNPs were formed in NEB Buffer 2.1 for 10 min at room temperature. Substrate DNA with a 5' FAM label was added and samples were incubated for 5 min at room temperature. Aliquots were taken from the reactions at 1, 2, and 5 min and were quenched by adding EDTA. Samples were column purified before being analyzed by capillary electrophoresis. The area under the curve of peaks corresponding to cleaved DNA was divided by the area under the curve for peaks of cleaved plus uncleaved DNA to calculate the fraction of cleaved DNA.

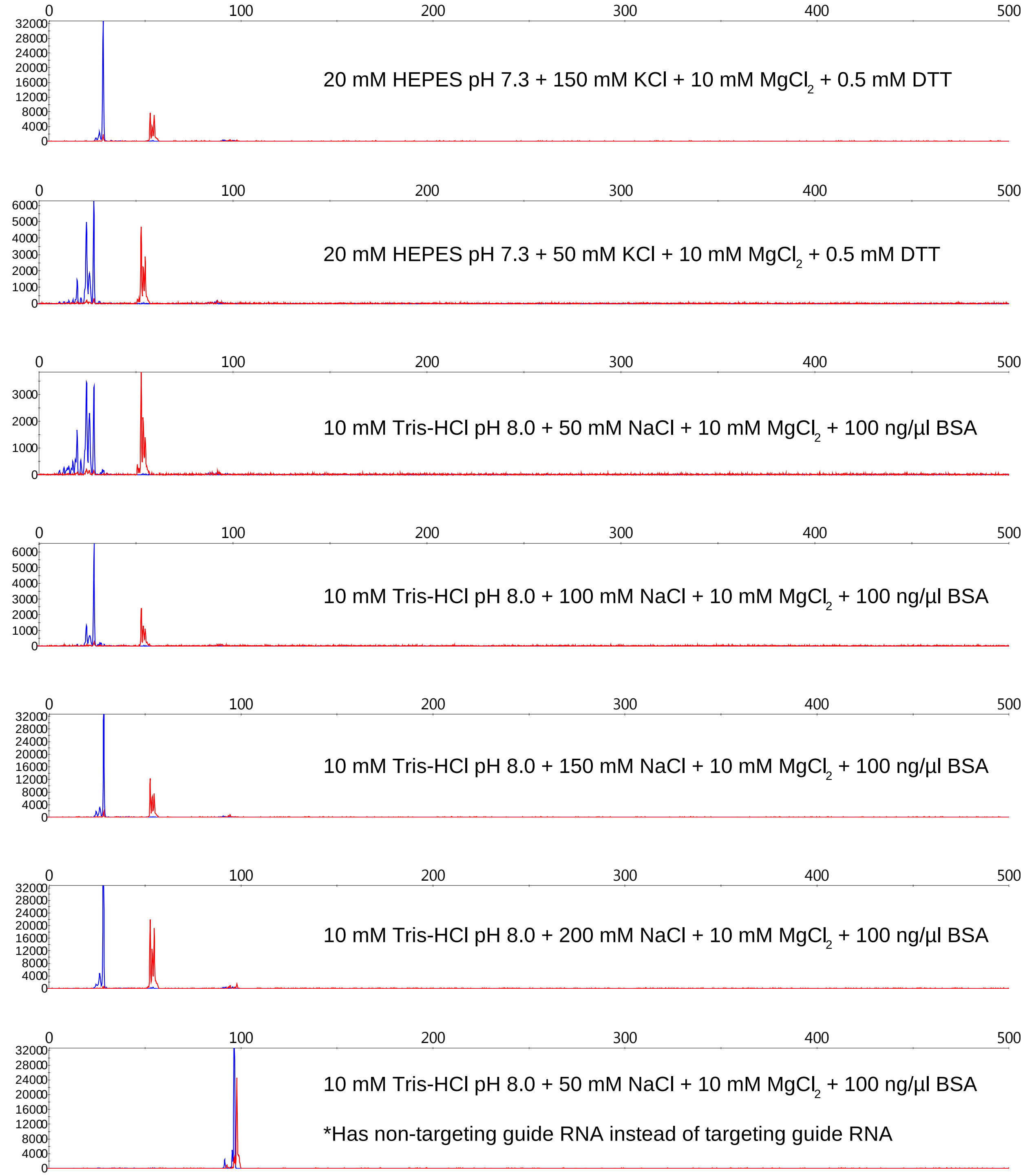

Supplementary Figure S5. Cleavage of on-target DNA activator by LbaCas12a in different reaction buffers. RNPs were formed in the indicated buffer composition for 10 min at room temperature. Substrate DNA with a 5’ FAM label on one strand and 5’ROX label on the other strand was added and samples were incubated for 5 min at 37°C. Reactions were quenched by adding EDTA and samples were column purified before being analyzed by capillary electrophoresis.

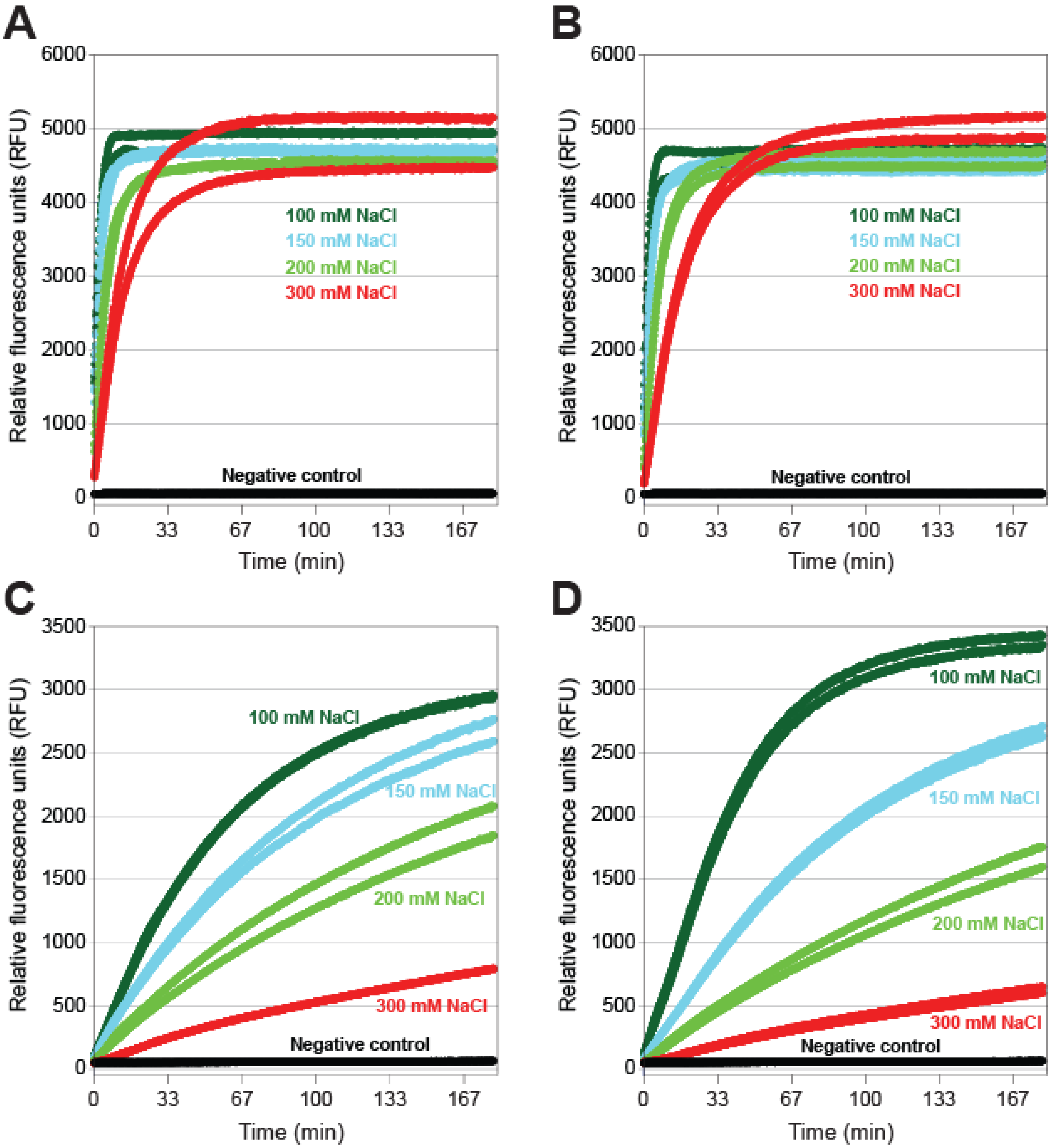

Supplementary Figure S6. Raw traces from fluorescent ssDNA and ssRNA reporter assays in different reaction buffers. All reaction buffers contained 10 mM Tris-HCl pH 7.9, 10 mM $MgCl_2$, 100 µg/µl BSA, and either 100 mM, 150 mM, 200 mM or 300 mM NaCl. **(A)** WTAP guide RNA, activator DNA, and fluorescent ssDNA reporter, **(B)** GFP2 guide RNA, activator DNA, and fluorescent ssDNA reporter, **(C)** WTAP guide RNA, activator DNA, and fluorescent ssRNA reporter, **(D)** GFP2 guide RNA, activator DNA, and fluorescent ssRNA reporter. In all cases, the negative control samples contained buffer with 100 mM NaCl and a guide RNA that did not target the included activator DNA.

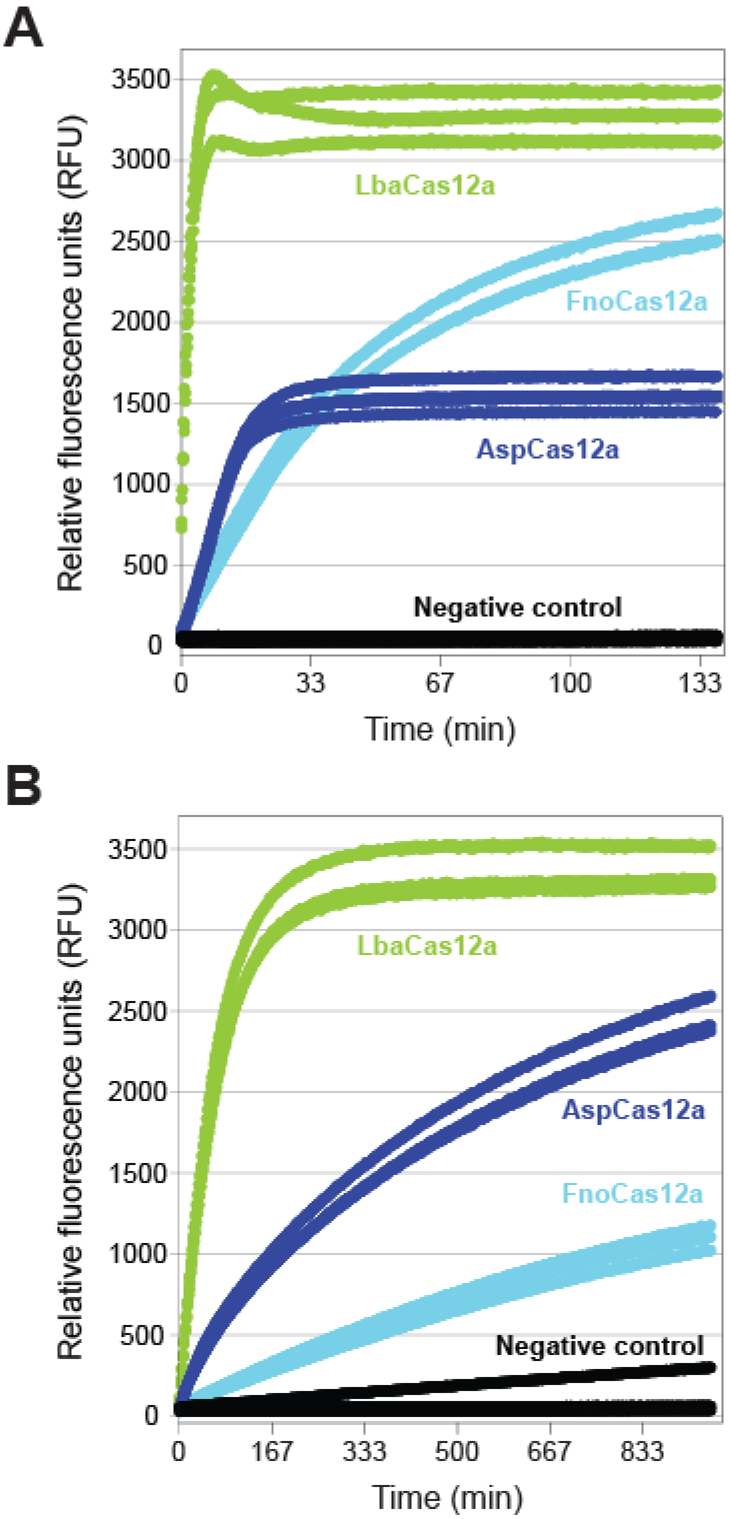

Supplementary Figure S7. Representative raw traces from fluorescent **(A)** ssDNA and **(B)** ssRNA reporter assays with different Cas12a orthologs. Reactions were carried out in NEB Buffer 2.1 for Lba and FnoCas12a. For AspCas12a a custom reaction buffer was used that consisted of 10 mM Tris-HCl pH 6.5, 10 mM $MgCl_2$, 100 mM NaCl, and 1 mM DTT. The negative control samples contained NEB Buffer 2.1 without any Cas protein added.

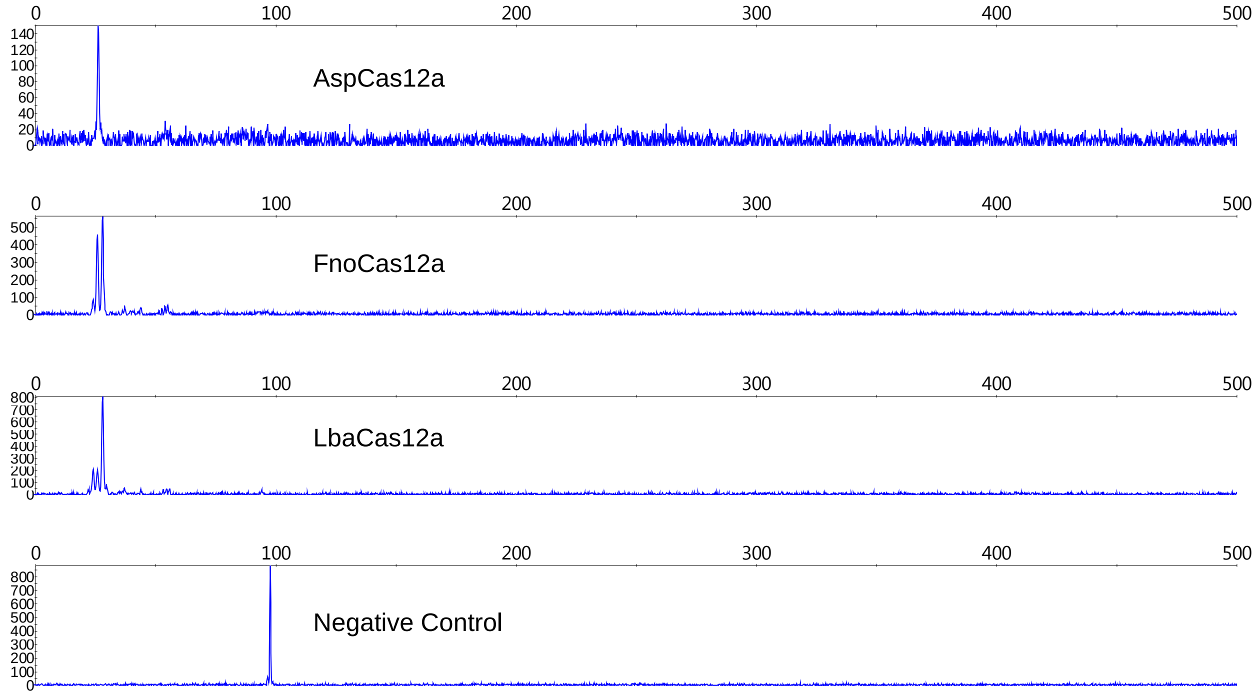

Supplementary Figure S8. Cleavage of on-target 5’ fluorescein-labeled DNA activator by Asp, Fno and LbaCas12a. Reactions were carried out in NEB Buffer 2.1 for Lba and FnoCas12a. For AspCas12a a custom reaction buffer was used that consisted of 10 mM Tris-HCl pH 6.5, 10 mM $MgCl_2$, 100 mM NaCl, and 1 mM DTT. Reactions were incubated for 10 min at 37°C, quenched with EDTA and purified using a Monarch PCR&DNA cleanup kit prior to analysis by capillary electrophoresis.
